# Varietal Reponses of rice (*Oryza sativa L*.) to different Nitrogen levels under Conventional Production Systems

**DOI:** 10.1101/2025.10.10.681668

**Authors:** Boadu Sober Ernest, Esther Fobi Donkor, Charles Afriyie-Debrah, Maxwell Darko Asante, Ralph K. Bam, Priscilla Francisco Ribeiro, Kirpal Agyemang Ofosu, Samuel Novo, Daniel Dzorkpe Gamenyah, Vincent Opoku Agyemang

## Abstract

Rice (Oryza sativa L.) is a vital staple crop globally, with its cultivation expanding to meet increasing demand. In Sub-Saharan Africa, particularly Ghana, rice productivity is often limited by poor soil fertility. Farmers frequently apply high nitrogen (N) fertilizer rates to boost yields; however, excessive nitrogen use contributes to environmental problems such as nutrient leaching and pollution. While optimal nitrogen application rates have been extensively studied, limited research has focused on varietal responses among rice genotypes. This study evaluated the response of five rice varieties (CRI-Agra Rice, Togo Marshall, CRI-Amankwatia, CRI-Enapa, and Jasmine 85) to different nitrogen rates (0, 30, 60, and 90 kg N/ha), focusing on morphophysiological, biomass and yield-related traits.

The findings showed significant (p < 0.001) variations in these traits with increasing nitrogen levels. Application of 90 kg N/ha led to substantial improvements: 40% increase in chlorophyll content, 34.3% in culm length, 71.4% in panicle number, 28.3% in straw dry weight and 42.9% in grain yield over the control (0 kg N/ha). Nitrogen significantly promoted vegetative growth, delayed flowering and enhanced biomass and grain production. Genotypic differences in nitrogen use efficiency were also observed. Togo Marshall, CRI-Agra Rice and Jasmine 85 showed over 30% increases in chlorophyll content, while CRI-Enapa exhibited higher plant height and panicle number at 90 kg N/ha. Togo Marshall and CRI-Enapa recorded the highest biomass and yield responses, indicating superior nitrogen utilization.

Overall, CRI-Enapa and Togo Marshall performed best at 60–90 kg N/ha. These findings highlight the importance of genotype-specific nitrogen management strategies for improving rice productivity and sustainability in Ghana and similar regions.

## 1.0 Introduction

Rice is a major staple food crop, feeding more than half of the global population. Its production is inherently nutrient-intensive: nitrogen (N) is a critical macronutrient driving photosynthesis and grain filling. In low-fertility soils, as often found in Sub-Saharan Africa (SSA), rice growth is strongly limited by N deficiency (Zingore *et al*., 2022[1] and Tsujimoto, 2025[2]).

Globally, farmers have applied more N fertilizer to increase yields, but this has led to environmental pollution and in efficiencies (Srikanth *et al.,* 2023[3]). On average, world fertilizer use is ∼146 kg/ha, whereas in Sub-Saharan Africa it is only ∼22 kg/ha. The low input rates in Africa contribute to persistent yield gaps: average rice yield in Africa (∼2.4 t/ha) is far below that in Asia (∼4.3 t/ha). This gap is attributed in part to insufficient nutrient supply and less efficient agronomy in African systems.

In Ghana, rice is widely cultivated under rain fed conditions, but average yields remain low (1–3 t/ha) compared to potential. National guidelines recommend N application rates of about 60–90 kg/ha for target yields of 3–4 t/ha (6, 3), yet farmers often apply less due to cost or knowledge gaps. Adequate N fertilization is known to enhance rice growth (tiller formation, leaf area, chlorophyll content) and yield components (panicle number, grain set) (Wang *et al*., 2022[4] and Srikanth *et al.,* 2023[3]). However, excess N beyond the crop’s needs can lower nitrogen use efficiency (NUE) and even reduce yields due to lodging or delayed maturity (Wang *et al.,* 2022[4] and Srikanth *et al.,* 2023[3]). Therefore, it is crucial to identify the optimal N rates that maximize yield without waste.

Importantly, rice genotypes vary in their response to N. High-yielding modern varieties often require high N inputs, while some local landraces or improved lines can maintain yield under lower N (Srikanth *et al.* (2023[3]). Studies in Asia have documented genotypic variability for traits such as N uptake and utilization efficiency, with strong links between NUE and grain yield (Opuni *et al*., 2023[5] and Srikanth *et al.,* 2023[3]). According to Srikanth *et al.* (2023)[3] rice yields increased significantly with moderate N rates. However, there is a dearth of information on genotype-specific N responses in West African rice germplasm, including Ghana (Opuni *et al*., 2023[5]). This knowledge gap hampers precise fertilization recommendations.

This study thus evaluates the responses of five rice genotypes to different N fertilizer rates for growth and yield parameters under conventional farming system in Ghana. The study aimed to determine optimal N rates for each variety and assess genotype × N interactions. The results were interpreted in light of comparable research on N response and NUE, to inform genotype-specific fertilization strategies that improve rice productivity and resource use efficiency in Ghana.

## 2.0 Materials and Methods

### 2.1 Study area

Two independent field experimentation was carried out in the major (March, 2023 - July, 2023) and minor (September - December, 2023) growing season at CSIR-Crops Research Institute, Rice Breeding Fields (6°42’4.0728"N, 1°31’53.364"W) in Fumesua, Ejisu municipality, which falls within the Forest agro-ecological zone in Ghana. Annual temperatures range from a minimum of 21.1°C to a maximum of 32.7°C and a mean of 31.6°C (Figure 3.1). The average annual rainfall is 1550 mm, and the seasonal distribution of rainfall is uneven; there is less precipitation in the third quarter of the year (Agyemang *et al*., 2023[6]). Figure 1 shows the map of the research site.

**Figure 1:**
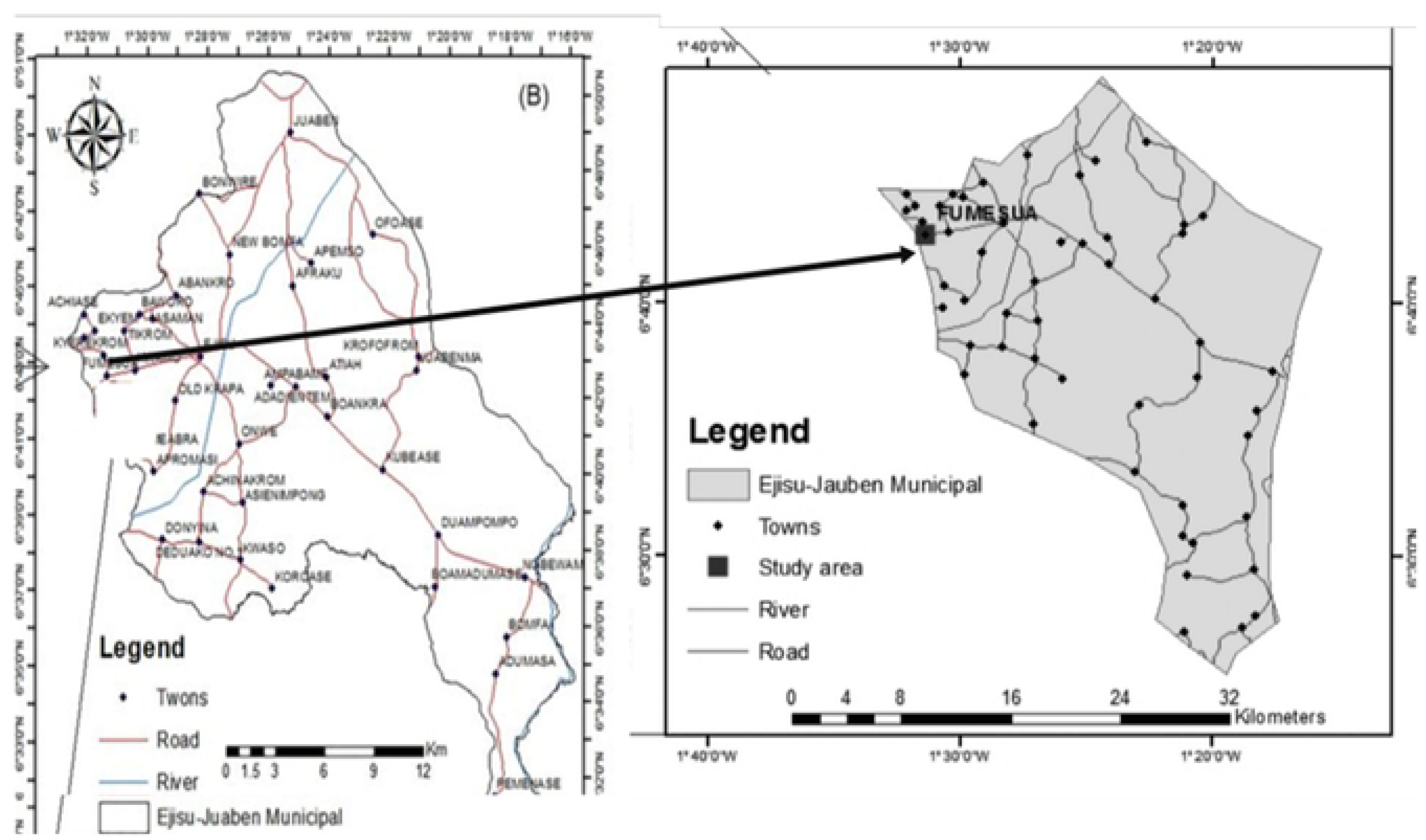
Map showing Ejisu-Juaben Municipal and CSIR-Crops Research Institute, Fumesua

### 2.2 Plant Materials

The research used five (5) rice varieties (Togo Marshall, CRI-Amankwatia, CRI-Agra rice, Jasmine 85 and CRI-Enapa) which are adapted to low-input conditions. These varieties consisted 4 released varieties from the Crop Research Institute, Council for Scientific and Industrial Research (CSIR-CRI, Fumesua-Kumasi). Table 1 shows the characteristics of the genetic materials used for the study.

**Table 1:** Characteristics of genetics materials used for the study

| Varieties | Year of release | Maturity Period (days) | Attribute | Yield Potential (t/ha) |
| --- | --- | --- | --- | --- |
| CRI-Agra Rice | 2013 | 125-130 | Long, slender<br>white | 7.5 - 8.0 |
| Togo Marshall | - | 120-125 | Long, slender<br>white | 7.0 - 7.5 |
| CRI-Amankwatia | 2010 | 115-120 | Long, slender<br>white | 7.5 - 8.0 |
| Jasmine 85 | 2013 | 120-125 | Long, slender<br>white | 6.5 - 7.0 |
| CRI-Enapa | 2017 | 130-135 | Long, slender<br>white and<br>aromatic | 8.0 - 8.5 |

### 2.3 Experimental Design and Field Layout

The experiment in both seasons was laid out in randomized complete block design in a split-plot arrangement with three replications. The trial consisted two experimental factors which were five (5) rice varieties (Togo Marshall, CRI-Amankwatia, CRI-Agra rice, Jasmine 85 and CRI-Enapa) and four (4) application rates of nitrogen fertilizer (0, 30, 60 and 90 kg N ha^−1^) The main plot treatments were the five (5) rice varieties while the sub-plot treatments were the four levels of nitrogen fertilizer.

Plot size was 4 m × 5 m, with 20 cm row spacing and seedlings planted at 20 cm × 20 cm spacing. The N rates were chosen to encompass the recommended range for target yields up to 4–5 t/ha (APNI Rice Cropping Guide (2021[7]), plus a zero-N control. Prior to field demarcation, the field was rotovated, levelled and then allowed to stay (lay fallow) for two weeks for volunteer crops to grow together with weeds. The weeds and volunteer crops were then controlled by spraying with Roundup before being manually hoed to prepare the plots prior to transplanting.

### 2.4 Soil characteristics

Before transplanting, soil samples were collected at different locations for compositional analysis at the CSIR-Soil Research Institute, Soil and Plant Chemistry laboratory, Kwadaso. The initial physicochemical properties of experimental soil are presented in Table 2.

**Table 2:**
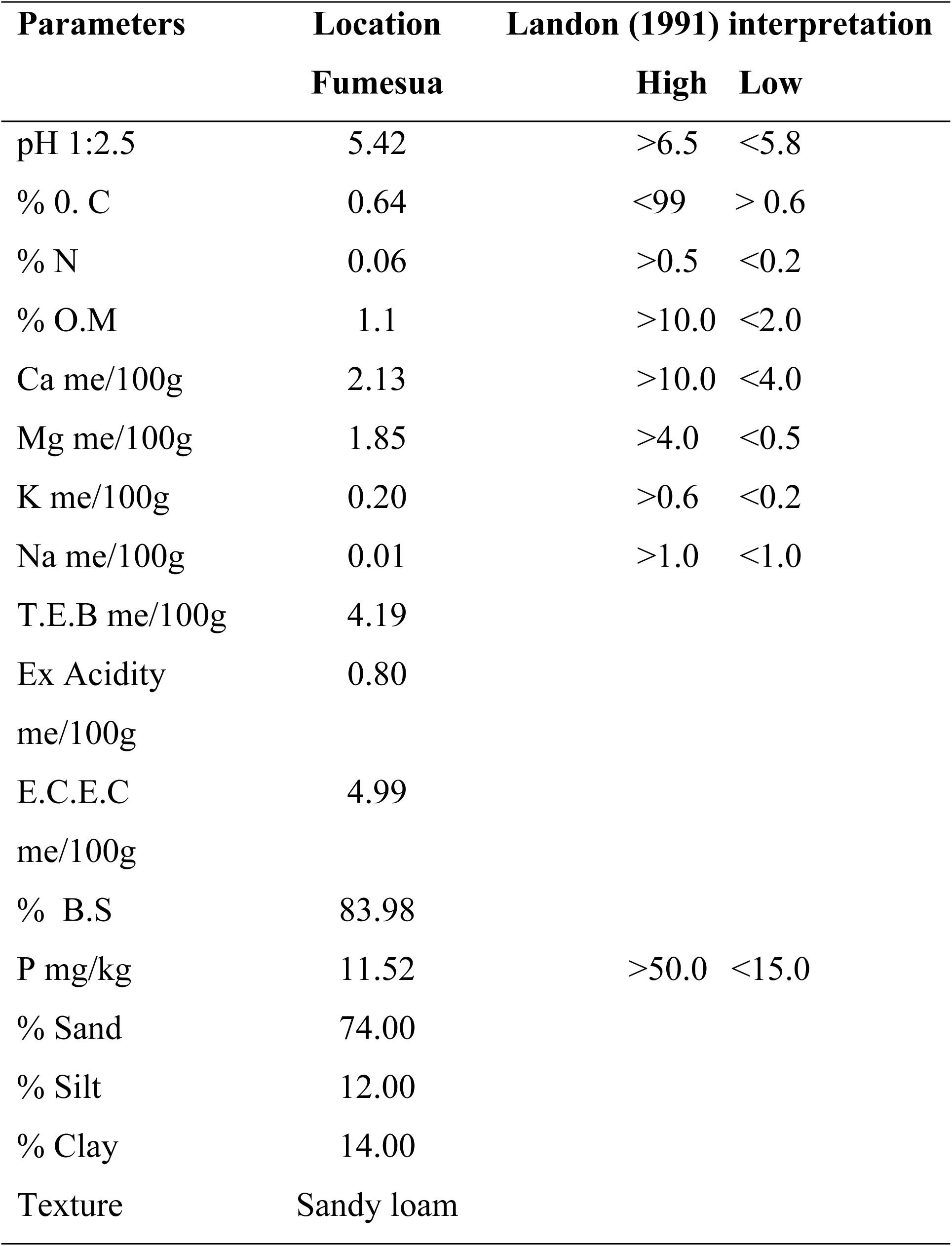
Initial Physio-chemical and mineralogical characteristics of soil samples from experimental sites

| Parameters | Location<br>Fumesua | Landon (1991) interpretation |  |
| --- | --- | --- | --- |
|  |  | High | Low |
| pH 1:2.5 | 5.42 | >6.5 | <5.8 |
| % 0. C | 0.64 | <99 | > 0.6 |
| % N | 0.06 | >0.5 | <0.2 |
| % O.M | 1.1 | >10.0 | <2.0 |
| Ca me/100g | 2.13 | >10.0 | <4.0 |
| Mg me/100g | 1.85 | >4.0 | <0.5 |
| K me/100g | 0.20 | >0.6 | <0.2 |
| Na me/100g | 0.01 | >1.0 | <1.0 |
| T.E.B me/100g | 4.19 |  |  |
| Ex Acidity<br>me/100g | 0.80 |  |  |
| E.C.E.C<br>me/100g | 4.99 |  |  |
| % B.S | 83.98 |  |  |
| P mg/kg | 11.52 | >50.0 | <15.0 |
| % Sand | 74.00 |  |  |
| % Silt | 12.00 |  |  |
| % Clay | 14.00 |  |  |
| Texture | Sandy loam |  |  |

### 2.5 Environmental condition during the study period

During the major season of the experiment, average temperatures ranged from 24.2°C in July to 27.2°C in March. Relative humidity was generally high, peaking in June at 86%, which coincided with the highest rainfall of 157 mm. Rainfall during this period was consistently substantial, with May (128 mm) and July (125 mm) also receiving notable amounts (Figure 2).

**Figure 2:**
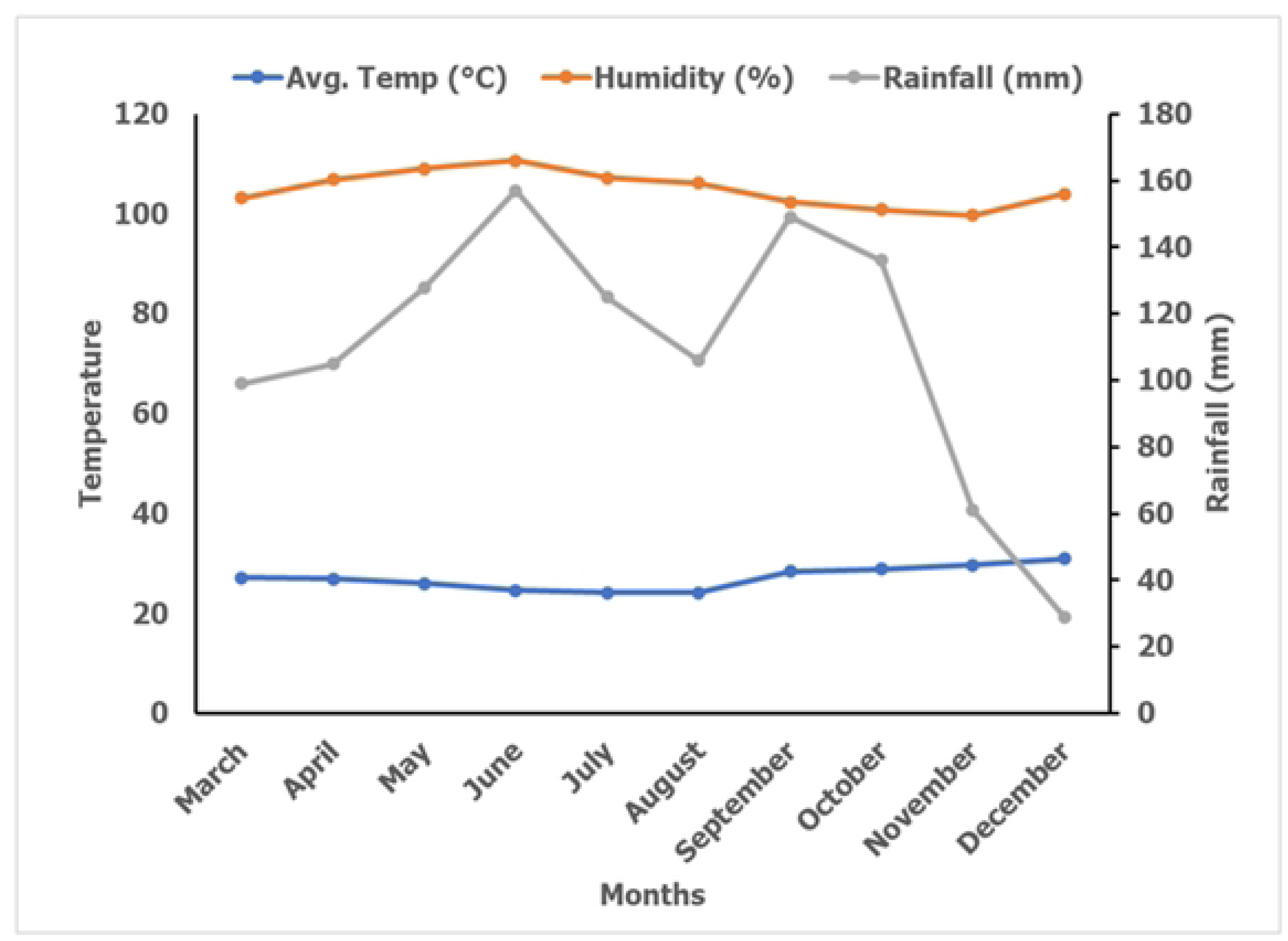
Climatic condition of the experimental field during the study period.

**Figure 3:**
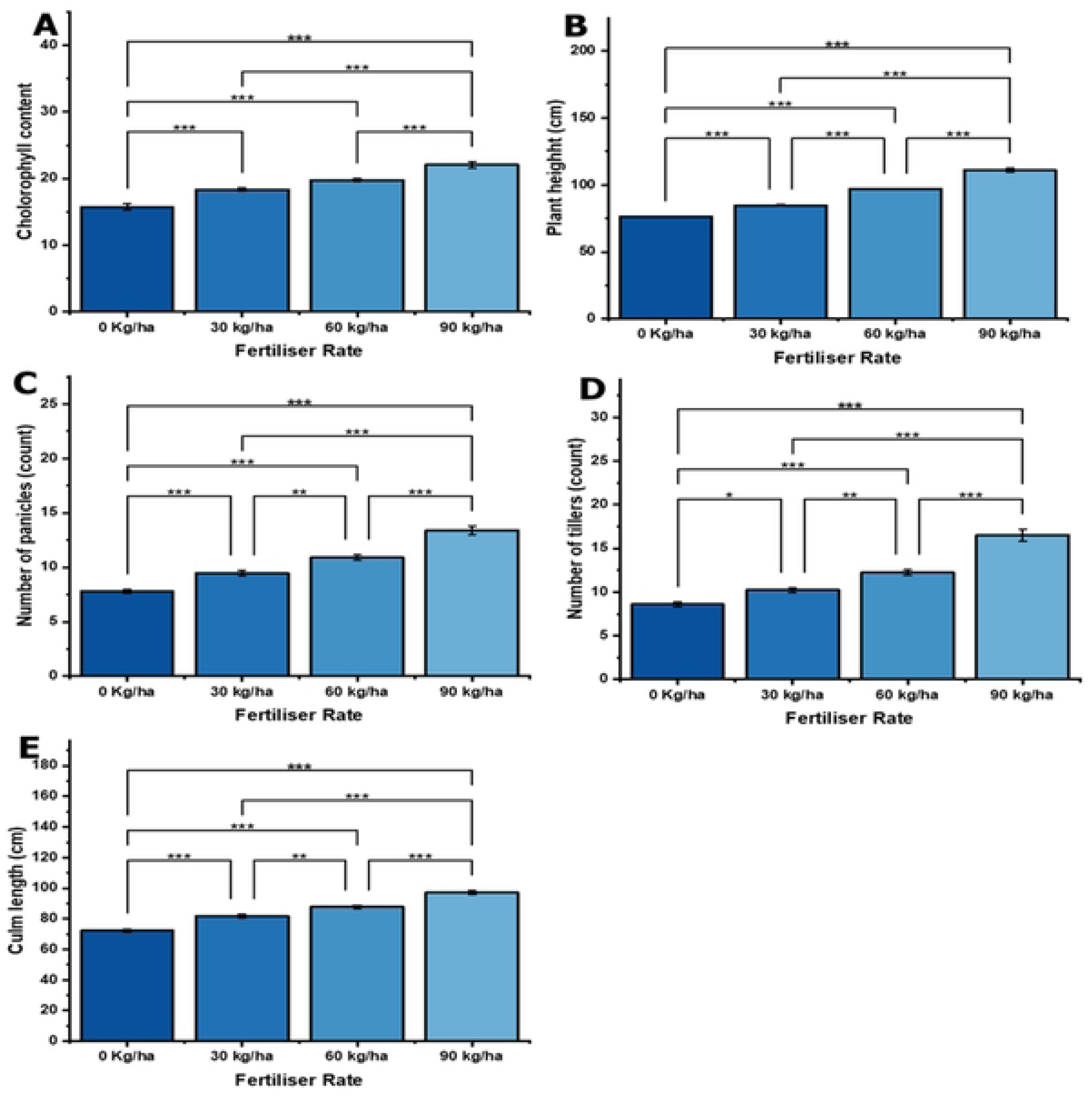
Effect of different nitrogen application rate on (A) chlorophyll content; (B) Plant height; (C) Number of panicles; (D) Number of tillers and (E) Culm length of rice. Where, error bar represents standard error of mean (SEM). Asterisks indicate significance levels: * p<0.05; P < 0.05P<0.05, ** p<0.01 and*** p<0.001.

In the minor season of the experiment, temperatures were higher, ranging from 28.4°C in September to 31.0°C in December. This period recorded lower humidity levels, ranging from 70% in November to 74% in September. Rainfall was more variable, with September still receiving a high amount (149 mm), but dropping significantly to 29 mm in December (Figure 2).

### 2.6 Nursery, transplanting and cultural practices

To establish rice seedlings for the study, a nursery was set up. A nursery bowl (0.8 m × 0.5 m) filled with topsoil dug at 0 – 15 cm depth from soil surface was used for establishing nursery for rice varieties. For each rice variety, healthy uniform seeds were pregerminated by soaking in water for 24 hours. Subsequently, the water was removed and the seeds were placed in a shaded area for two days. The nursery bowl filled with soil was then sown with these pre-germinated rice seeds. After a period of 21 days, healthy and uniform growing rice seedlings of each variety were transplanted to the demarcated experimental field using a planting distance of 20 cm × 20 cm. Both experiments were rain-fed, however irrigation was augmented with irrigation systems as and when needed. Weeds in the experimental fields were controlled by hand picking as well as application of selective weedicides.

The experimental fields were sprayed with a systemic insecticide K-Optimal (Lambda Cyhalothrine 15 g/L +Acetamipride 20 g/L EC) to control common rice pests. All experimental plots were covered with a net at the flowering stage of the rice to help prevent birds from feeding on the rice grains. The rice plant in the experiment area did not show any incidence of diseases and as such no disease control measure was employed.

### 2.7 Fertiliser rates imposition

Imposition of fertiliser treatment was carried out 2-weeks after transplanting to ensure uniform establishment of seedlings. Application of fertiliser was carried out using split application method thus, applying two halves separately at different times as shown in Table 3. The amount of chemical nitrogen applied to achieved each application rate was estimated based on Equation 1 below;

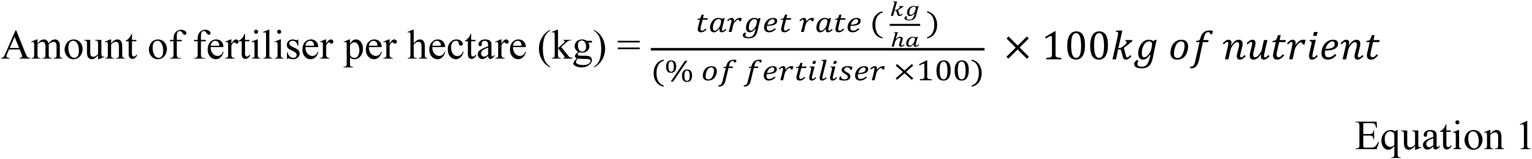

**Table 3:** Application rate of various nitrogen fertilisation levels used in the present study

| Nitrogen rate | Basal application (NPK 15:15:15) | Top dressing (Urea) |
| --- | --- | --- |
| Control (0 kg/ha) | 0 kg | 0 kg |
| 90 kg/ha | 45 kg | 45 kg |
| 60 kg/ha | 30 kg | 30 kg |
| 30 kg/ha | 15 kg | 15 kg |

### 2.8 Data Collection

Plant height and tiller number (productive tillers/m^2^) were recorded at flowering for 10 randomly selected hills per plot. Leaf chlorophyll content (SPAD values) was measured at mid-tillering. At maturity, each plot was harvested for yield and biomass data. Ten plants per plot were cut to determine number of panicles per plant, culm length and 1000-grain weight (adjusted to 14% moisture). Grain yield/plot was determined by threshing all panicles in a subplot, weighing the grain and converting to t/ha. Straw biomass (fresh straw weight and dry straw weight) was also measured.

### 2.9 Statistical Analysis

Data collected were entered into excel sheets and subjected to analysis of variance (ANOVA) using R statistical software package. Tukey’s test of significance (LSD; p = 95%) was used for mean separation among experimental variables, Statistical analysis was performed using model in Equation 2.

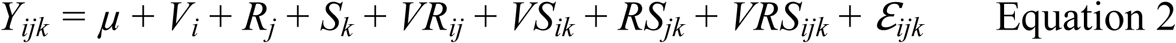

Where:

*Y_ijk_* represents the observation from *ijk^th^* rice variety, nitrogen rate and season, and
*μ* is the overall mean;
*V_i_ is* the effect of the *i^th^* variety;
*R_j_* is the effect of the *j^th^* nitrogen rate;
*S_k_* is the effect of the *k^th^* season of experiment;
*VR_ij_* is the interactive effect of the *i^th^* variety with *j^th^* nitrogen rate,
*VS_ik_* is the interactive effect of the *i^th^* variety with *k^th^* season,
*RS_jk_* is the interactive effect of the *j^th^* nitrogen rate with *k^th^* season,
*VRS_ijk_* is the interactive effect of the *i^th^* variety with *j^th^* nitrogen rate and *k^th^* season of experiment and
*_ijkl_* is the experimental error.

Person correlation analysis was carried using R (R Core Team, 2021[8]) to establish the relationship and association between measured parameters among rice varieties. Corrplot in R package version 0.84 (Wei et al, 2017[9]) was used to visualize the relationships among traits.

## 3.0 Results

### 3.1 Variability among the treatment combinations

The ANOVA analysis revealed significant differences (*p< 0.001*) among the five (5) rice varieties for all the traits studied (Table 4). Nitrogen application was also significant (*p < 0.001*) for all the traits measured (Table 4). The analysis also revealed significant season effect on number of panicles (*p = 0.004*), chlorophyll content (*p = 0.001*), straw fresh weight (*p = 0.003*) and number of tillers (*p < 0.001*). However, plant height, culm length, days to flowering, straw dry weight, 1000-grain weight, and yield showed non-significant(*p>0.05*) season effect. (Table 4).

**Table 4.**
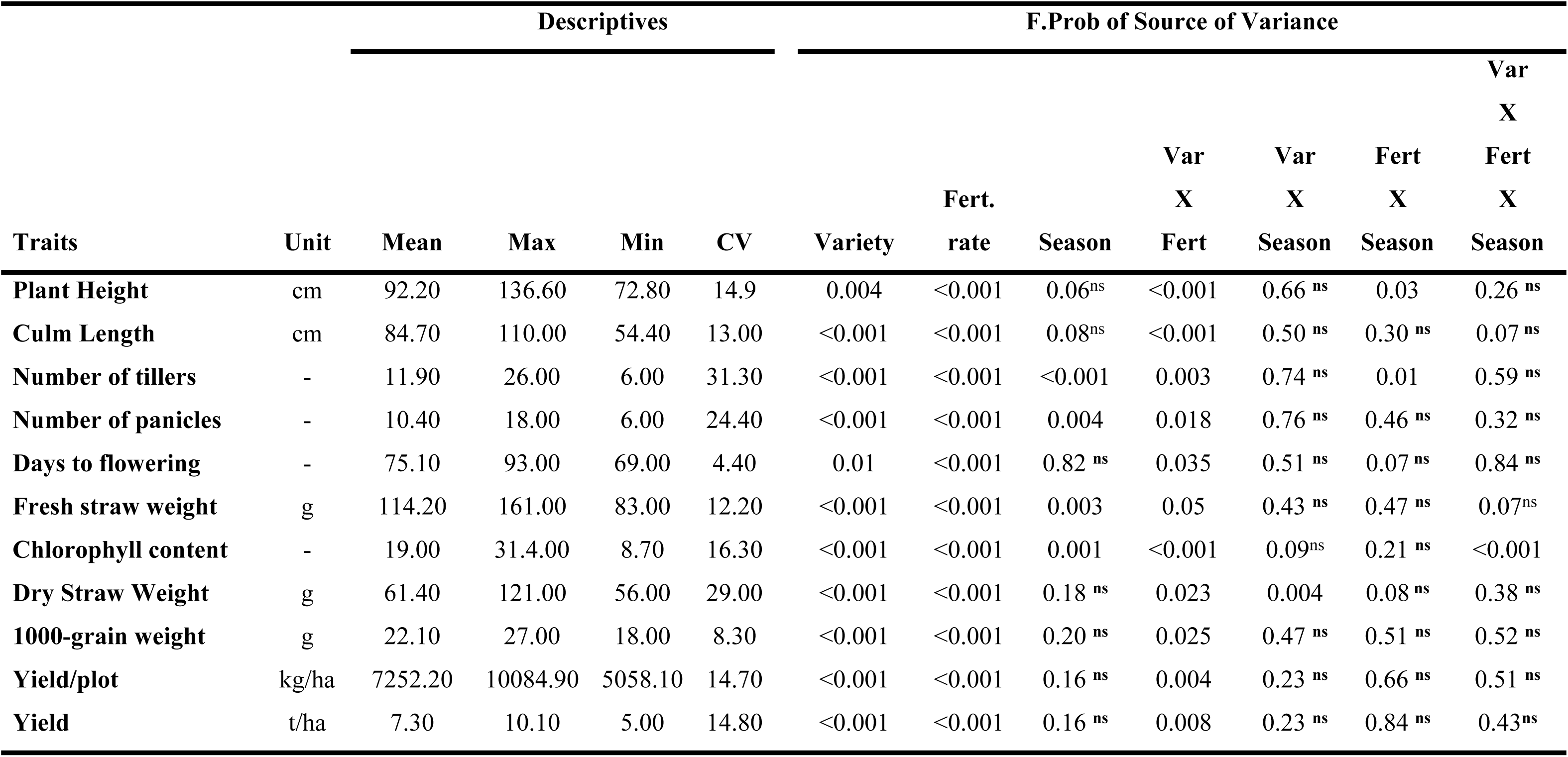
Summary Analysis of Variance (ANOVA) of measured morphological, yield and biomass parameters of rice varieties cultivated under different N fertilization.

Interaction of variety and fertiliser rate was significant (*p < 0.05*) all the traits studied with plant height, culm length and chlorophyll content recording high significant levels (*p<0.001*) (Table 4).

The results also showed that interaction of variety and season was non-significant (*p > 0.05*) for plant height, 1000-grain weight, number of tillers, number of panicles and yield. However, there was significant interaction of variety and season for straw dry weight (*p = 0.004*) (Table 4). Except for number of tillers (*p = 0.01*) and plant height (*p = 0.025*), interaction of fertiliser rate and season was non-significant (*p >0.05*). The analysis also revealed non-significant (*p>0.05*) effect of the three-way interaction of variety, fertiliser rate and season except for chlorophyll content (*p < 0.001*) (Table 4).

### 3.2 Effect of fertiliser rate on growth parameters

Fertiliser application significantly affected measured morphophysiological traits (Figure 3 A-E). About 1.4-fold variation in chlorophyll content was observed between the upper and lower boundaries of fertiliser rates (Figure 3A). Chlorophyll content showed a direct association with increasing fertiliser rate. When cultivated under 90 kg N/ha (22.1), 11.5%, 20% and 40.1% increase in chlorophyll content was recorded compared to 60 kg N/ha (19.8), 30 kg N/ha (18.4) and 0 kg N/ha (15.8) respectively. Compare to the control (zero N application), 60 kg N/ha and 30 kg N/ha had 25.6% and 16.5% increase in chlorophyll content (Figure 3A).

Similarly, plant height varied 1.6-fold among various fertiliser rates (Figure 3B). Application of nitrogen at 90 kg N/ha resulted in 14.5% - 45.1% increase in plant height compared to 60 kg N/ha, 30 kg N/ha and 0 kg N/ha. When rice was cultivated under nitrogen unamended soil (control), 26.8% and 10.7% decrease in plant height was recorded compared to 60 kg N/ha and 30 kg N/ha respectively (Figure 3B). Compared to 30 kg N/ha, plant height was 14.5% higher under 60 kg N/ha (Figure 3B).

Number of panicles was relatively higher for 90 kg N/ha (13 count) as compared to 60 kg N/ha and 30 kg N/ha which recorded 10.9 and 9.5 count of panicles respectively (Figure 3C). On the other hand, control recorded the lowest number of panicles of 7.8 count (Figure 3C). As illustrated in Figure 3D, number of tillers varied 1.9-fold among fertiliser rates. In general, a direct association was observed between increasing fertiliser application and number of tillers. When grown under 90 kg N/ha, 35.2 – 92.2% increase in number of tillers was recorded as compared to 60 kg N/ha, 30 kg N/ha and 0 kg N/ha (Figure 3D). Compared to control, 60 kg N/ha and 30 kg N/ha had 42.2 and 18.9% increase in number of tillers respectively (Figure 3D).

Culm length responded positively to fertiliser application. Fertiliser application resulted in 1.3- fold variation in culm length among the rice varieties (Figure 3E). It was clear that, soil amended with nitrogen fertiliser recorded higher mean culm length compared to the control. Application rate at 90 kg N/ha recorded 34.3% increase in culm length compared to control whereas 60 kg N/ha and 30 kg N/ha recorded 21.4% and 13.3% increase in culm length respectively compared to control which had the lowest culm length of 72.3 cm (Figure 3E).

### 3.3 Interaction of variety and nitrogen application on growth parameters

Across all varieties, chlorophyll content increased positively in response to nitrogen application. However, the magnitude of increase varied significantly (p < 0.001) among varieties (Figure 4A - D). When cultivated at 90 kg N/ha, there was 26%, 30.4%, 36.1%, 38.9% and 81.7% increase in chlorophyll content for CRI-Enapa, CRI-Amankwatia, CRI-Agra, Togo Marshall and Jasmine 85 respectively (Figure 4A). Despite the observed genotypic variation in response to N-fertilisation, genotypes showed no significant variation in chlorophyll content when cultivated under 30 kg N/ha and 60 kg N/ha (Figure 4A).

**Figure 4:**
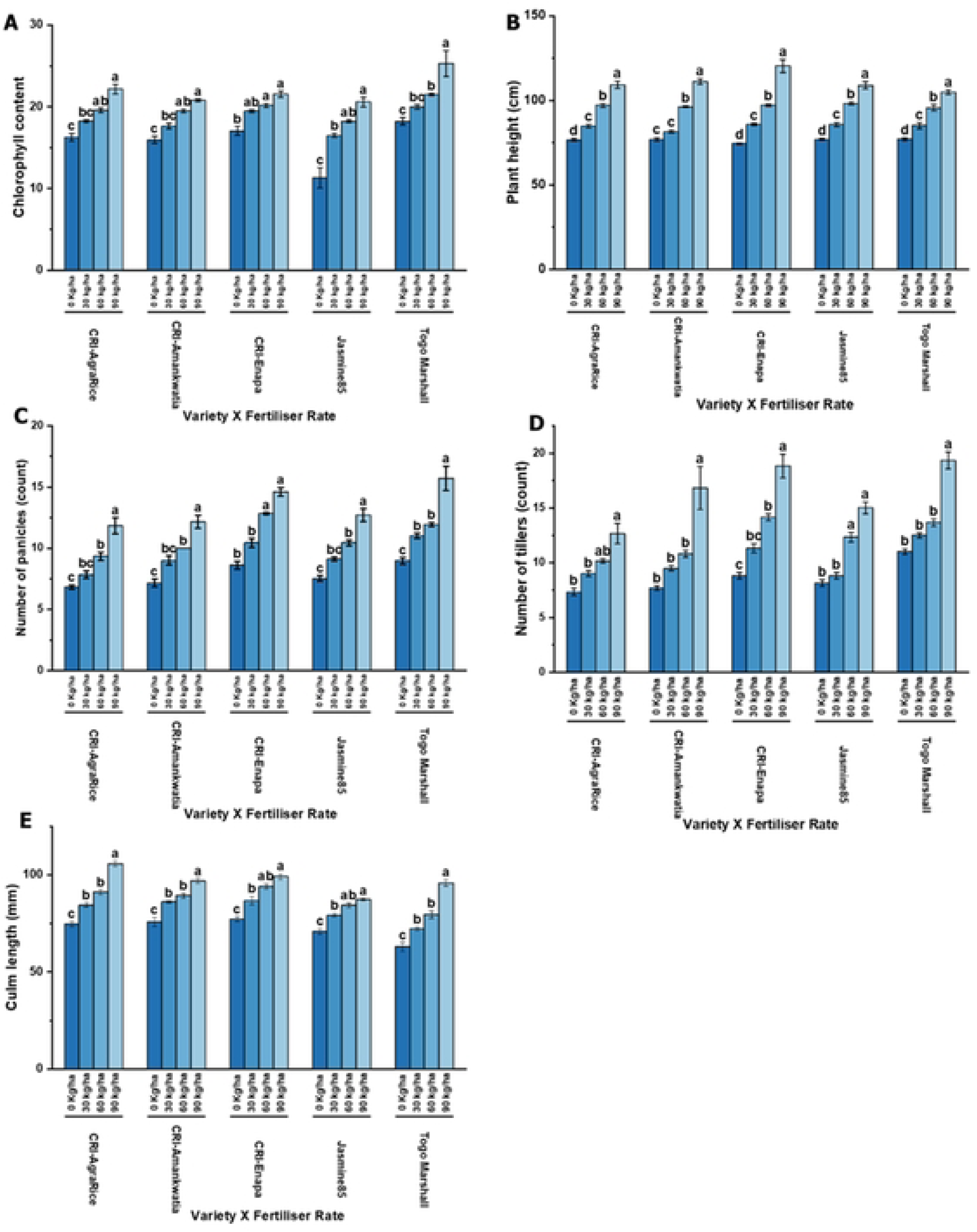
Interaction of varieties and nitrogen application rate on (A) chlorophyll content; (B) Plant height; (C) Number of panicles; (D) Number of tillers and (E) Culm length of rice. Where, error bar represents standard error of mean (SEM). Means with similar variable indicating non-significance while different variables indicate significance (p< 0.05 or 0.001).

Plant height among varieties increased with increasing nitrogen application (Figure 4B). Plant height was relatively lower at 0 kg N/ha however, CR1-Amaankwatia recorded statistically similar plant height at 0 kg N/ha (76.98 cm) and 30 kg N/ha (81.7 cm) (Figure 4B). The percentage increase in plant height of 61.7% was recorded for CRI-Enapa while 36.1 - 44.4% was recorded by Jasmine 85, Togo Marshall, CRI-Agra rice and CRI-Amankwatia under 90 kg N/ha application rate (Figure 4B).

The response of rice varieties to nitrogen application showed a consistent increase in number of panicles across all fertilizer rates (Figure 4C). Thus, number of panicles improved as nitrogen levels increased from 0 kg/ha to 90 kg/ha, though the magnitude of response varied among the different varieties (Figure 4C). Togo Marshall showed the highest number of panicles at all fertilizer rates, reaching 15.7 at 90 kg/ha, making it the most responsive variety to N application. CRI-Enapa also exhibited a strong response, increasing from 8.6 at 0 kg/ha to 14.6 at 90 kg/ha. Jasmine85 showed a moderate response, with yield increasing from 7.53 to 12.7. CRI-Agra rice and CRI-Amankwatia followed a similar trend but had relatively lower yields compared to CRI-Enapa and Togo Marshall (Figure 4C). Despite variation in number of panicles among varieties, panicles were statistically similar at 30 kg N/ha and 60 kg N/ha for Jasmine, Togo Marshall, CRI-Agra rice and CRI-Amankwatia (Figure 4C).

The direct response of number of tillers to nitrogen application was observed among rice varieties in the study (Figure 4D). The highest tiller number at 90 kg/ha was observed in Togo Marshall (19.3 tillers), followed by CRI-Enapa (18.8 tillers) and CRI-Amankwatia (16.8 tillers). CRI-Agra rice had the lowest number of tillers (12.7) at 90 kg/ha, indicating a comparatively lower response to nitrogen application. The increase in tiller count was most pronounced between 60 kg/ha and 90 kg/ha, particularly in CRI-Amankwatia and Togo Marshall (Figure 4D). At 0 kg N/ha, number of tillers was highest for Togo Marshall (11 tillers), followed by CRI-Enapa (8.83 tillers), while CRI-Agra Rice had the lowest (7.33 tillers). CRI-Amankwatia and Enapa displayed a sharp increase of over 100% at 90 kg/ha, suggesting strong tillering response to nitrogen (Figure 4D).

The response of culm length to nitrogen (N) application showed an increasing trend across all varieties (Figure 4E). Thus, culm length increased with increasing nitrogen application across all rice varieties, indicating a positive response to nitrogen fertilization. The highest culm length at 90 kg/ha was observed in CRI-Agra rice (105.8 cm), followed by CRI-Enapa (99.27 cm) and CRI-Amankwatia (96.97 cm). Togo Marshall, which had the shortest culm length at 0 kg/ha (62.87 cm), exhibited the greatest increase in length at 90 kg/ha (95.85 cm), suggesting a strong response to nitrogen application (Figure 4E). The lowest increase in culm length was observed in Jasmine 85, which reached only 87.33 cm at 90 kg/ha, indicating a relatively lower response to nitrogen compared to other varieties (Figure 4E).

### 3.4 Effect of nitrogen application on yield and yield contributing traits

The application of nitrogen (N) fertilizer influenced the number of days to flowering in rice, with an increasing trend observed as nitrogen levels increased (Figure 5A). Higher nitrogen rates delayed flowering across all treatments, as indicated by the increasing number of days to flowering with increasing nitrogen application. At 0 kg/ha, rice flowered the earliest at 71.63 days, while at 90 kg/ha, flowering was delayed to 78.93 days (Figure 5A). A steady increase in days to flowering was observed from 71.63 days (0 kg/ha) to 73.97 days (30 kg/ha), 75.90 days (60 kg/ha), and 78.93 days (90 kg/ha) (Figure 5A).

**Figure 5:**
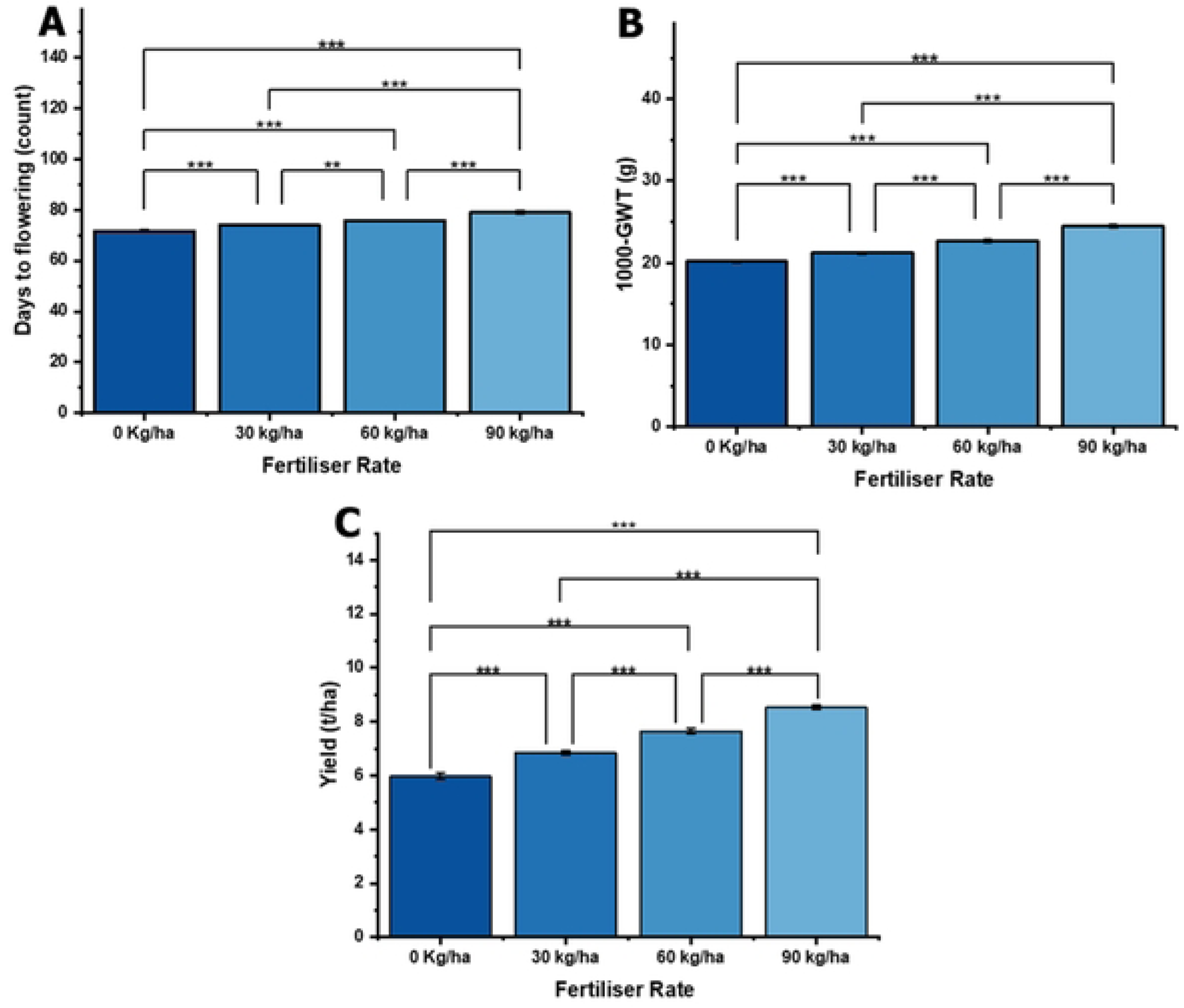
Effect of different nitrogen application rate on (A) Days to flowering; (B) 1000-grain weight and (C) Grain yield. Where, error bar represents standard error of mean (SEM). Asterisks indicate significance levels: *p <0.05; P < 0.05P<0.05, ** p<O.OI and*** p<O.001.

A direct relationship between nitrogen application and 1000-graain weight was observed as illustrated in Figure 5B. 1000-grain weight ranged from 20.2 g – 24.5 g for 0 kg N/ha and 90 kg N /ha respectively (Figure 5B). Compared to control, 5%, 112.4% and 211.3% increase in 1000-grian weight was recorded when rice was grown under 30 kg N/ha, 60 kg N/ha and 90 kg N/ha respectively (Figure 5B). 60 kg N/ha recorded 6.9% increase in 1000-grain weight compared to 30 kg N/ha (Figure 5B).

Yield increased progressively with increasing nitrogen rates, indicating a strong positive response of rice to nitrogen application. The lowest yield was observed at 0 kg/ha (5.97 t/ha), while the highest yield was recorded at 90 kg/ha (8.6 t/ha). A gradual yield increase was noted at intermediate nitrogen rates, with 6.9 t/ha at 30 kg/ha and 7.7 t/ha at 60 kg/ha (Figure 5C).

### 3.5 Interaction of variety and nitrogen application on yield and yield contributing parameters

The interaction of rice variety and nitrogen fertilizer application influenced the days to flowering. Generally, an increasing trend in the number of days to flowering was observed as nitrogen application rates increased across all varieties (Figure 6A). At 0 kg/ha, the days to flowering ranged from 70.67 days (Jasmine85 and Togo Marshall) to 73 days (CRI-AgraRice). As nitrogen application increased, all varieties exhibited delayed flowering, with the longest days to flowering recorded at 90 kg/ha, ranging from 77.17 days (CRI-Enapa) to 80.2 days (Togo Marshall). CRI-Agra Rice and CRI-Amankwatia showed a moderate delay in flowering, with 79 and 78.67 days at 90 kg/ha, respectively. Jasmine85 and Togo Marshall showed the most pronounced delay in flowering with 79.67 and 80.17 days at 90 kg/ha, respectively (Figure 6A).

**Figure 6:**
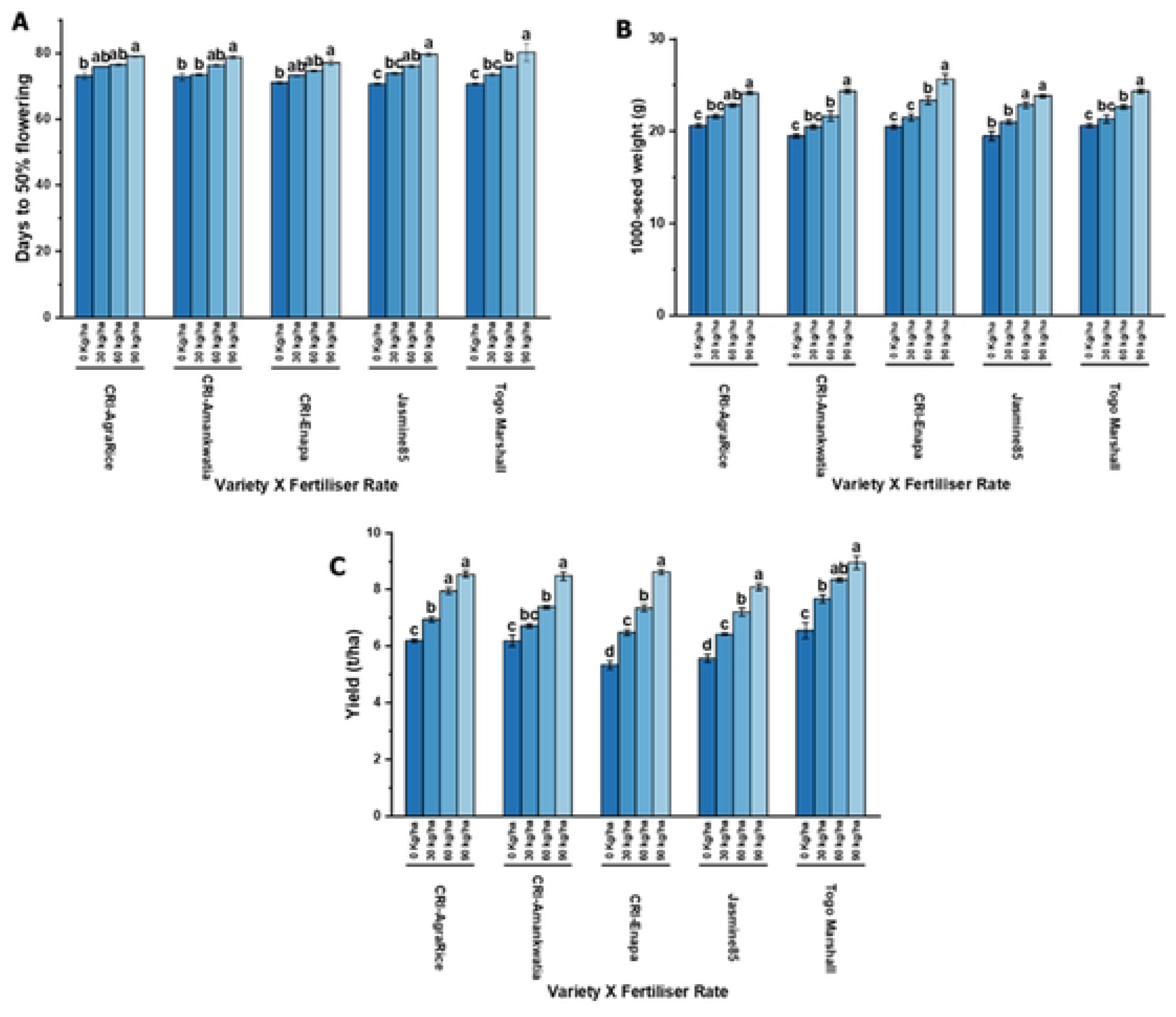
Interaction ofnitrogen and variety on (A) Days to flowering; (B) 1000-grain weight and (C) Grain yield. Where, error bar represents standard error of mean (SEM). Means with similar variable indicating non-significance while different variables indicate significance (p< 0.05 or 0.001).

The 1000-grain weight was influenced by both nitrogen application rates and variety (Figure 6B). A direct relationship between grain weight was observed with higher nitrogen levels across all varieties (Figure 6B). At 0 kg/ha, 1000-grain weight ranged from 19.5 g (CRI-Amankwatia, Jasmine85) to 20.67 g (CRI-AgraRice, Togo Marshall). As nitrogen application increased to 90 kg/ha, grain weight increased across all varieties, with values ranging from 23.83 g (Jasmine85) to 25.67 g (CRI-Enapa) (Figure 6B). The most significant improvement in grain weight was observed in CRI-Enapa, which increased from 20.5 g at 0 kg/ha to 25.67 g at 90 kg/ha. CRI-Amankwatia and Togo Marshall also showed considerable increases, reaching 24.33 g at 90 kg/ha. Jasmine85 had the lowest 1000-grain weight at all fertilizer levels but still showed a positive response to nitrogen application (Figure 6B).

The rice yield was significantly influenced by nitrogen application across all varieties. An increase in nitrogen levels resulted in a progressive improvement in yield, with notable variations among the different varieties (Figure 6C). At 0 kg/ha, the lowest yield was recorded in CRI-Enapa (5.3 t/ha), while the highest was observed in Togo Marshall (6.6 t/ha). Increasing nitrogen application to 90 kg/ha resulted in the highest yield for all varieties, with Togo Marshall achieving the highest yield (8.953 t/ha). CRI-Enapa showed the highest fold change (1.6) and the highest percentage increase (61.6%), indicating the strongest response to nitrogen fertilization. Jasmine85 exhibited a 44.98% increase (1.5-fold change), suggesting moderate nitrogen responsiveness. CRI-Agra rice, CRI-Amankwatia, and Togo Marshall had similar responses, with increases of approximately 37%, highlighting their relatively stable yield gains under higher nitrogen rates (Figure 6C).

### 3.6 Response of biomass production of rice varieties to nitrogen fertiliser application

Straw fresh weight ranged from 100.4 g – 1132.6 g indicating 1.3-fold variation among fertiliser application rates (Figure 7A). Compared to the control, 32%, 14% and 8.3% increase in straw fresh weight was recorded under 90 kg N/ha, 60 kg N/ha and 30 kg N/ha respectively (Figure 7A). Similarly, 90 kg N/ha nitrogen recorded 21.9% and 15.2% increase in straw fresh weight compared to 30 kg N/ha and 60 kg N/ha respectively (Figure 7A). Application of 60 kg N/ha (115.1 g) resulted in 5.9% increase in straw fresh weight compared to 30 kg N/ha (108.7 g) (Figure 7A).

**Figure 7:**
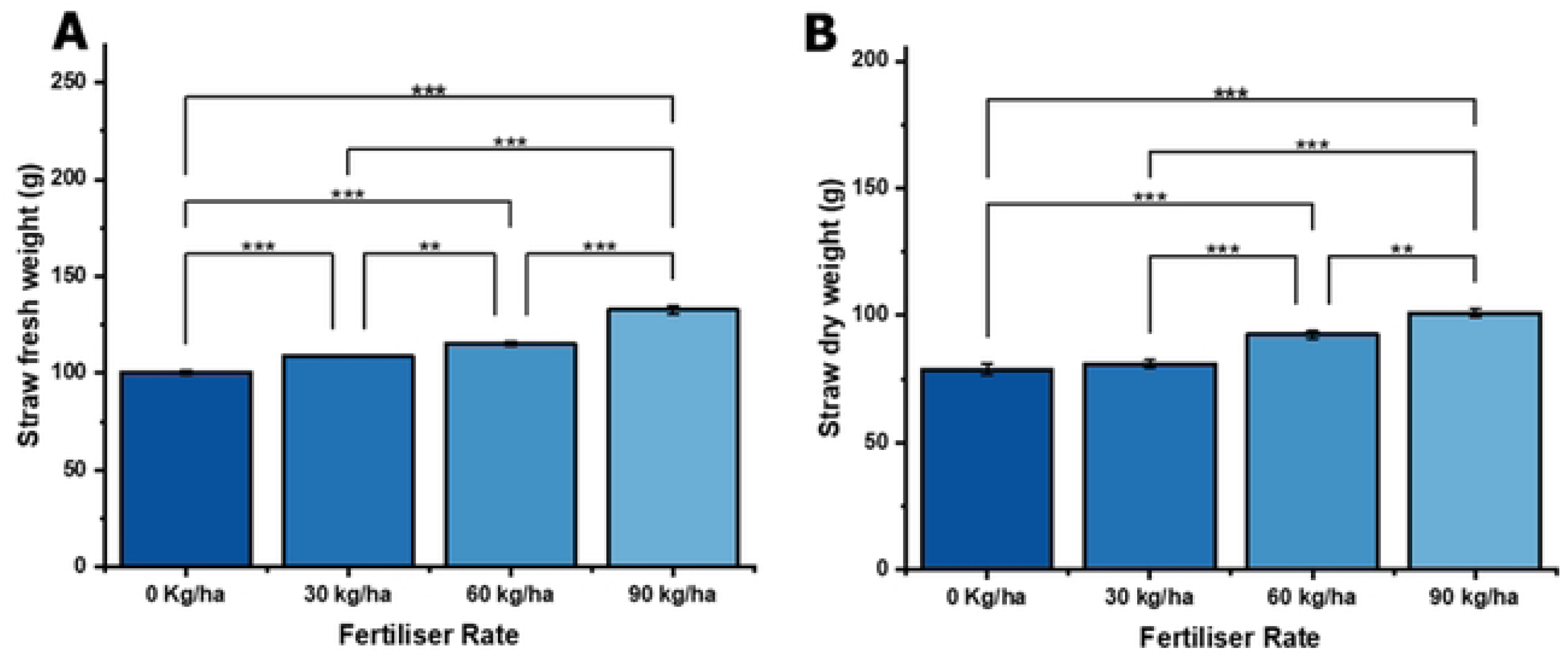
Effect of different nitrogen application rate on (A) Straw fresh weight and (B) Dry straw weight. Where, error bar represents standard error of mean (SEM). Asterisks indicate significance levels: * p<0.05; P < 0.05P<0.05, ** p<0.01 and*** p<0.001.

Dry straw weight exhibited 1.3-fold variation among fertiliser rates as illustrated in Figure 7B. Dry straw weight increased with increasing nitrogen rate with 90 kg N/ha recording the highest dry straw weight of 78.9 g (Figure 7B). Dry straw weight was 74 g and 75.9 g under 30 kg N/ha and 60 kg N/ha respectively (Figure 7B).

### 3.7 Interaction of variety and nitrogen application on biomass production

Rice varieties showed significant variation in their straw fresh weight in response to nitrogen application. However, the degree of response varied among varieties (Figure 8A-B). CRI-Agra rice and Jasmine 85 exhibited statistically similar responses (around 30% increase), while CRI-Amankwatia had the lowest increase (28.2%). At 90 kg N/ha, CRI-Enapa had the highest response to nitrogen fertilization, with a 37.02% increase, suggesting it has the greatest potential for biomass accumulation under high nitrogen levels. Similarly, Togo Marshall also showed a strong response (34.34%) increase indicating a substantial improvement in fresh weight with nitrogen application (Figure 8A). Despite 0 kg N/ha recording the lowest fresh straw weight across all varieties, fresh straw weight recorded at 0 kg N/ha, 30 kg N/ha and 60 kg N/ha were statistically similar for varieties CRI-Agra rice and CRI-Amankwatia (Figure 8A).

**Figure 8:**
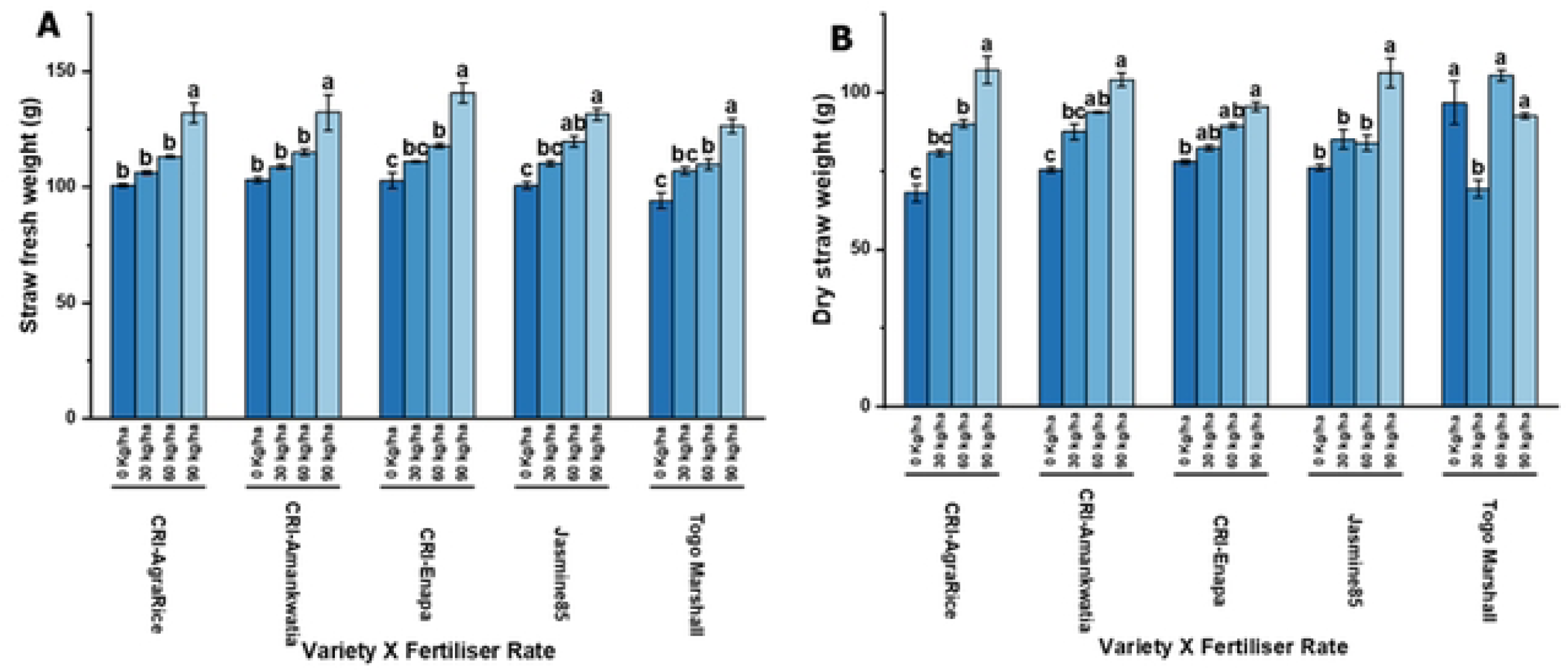
Interaction of variety and nitrogen application on biomass production. (A) straw fresh weight and (B) Straw dry weight. Where, error bar represents standard error of mean (SEM). Means with similar variable indicating non-significance while different variables indicate significance (p < 0.05 or 0.001).

Dry biomass of rice was significantly affected by nitrogen application rate (Figure 8B). In the present study, 0.1, 1.2, 1.4, 1.4 and 1.6-folds change in dry biomass was recorded between the upper and lower application rate for Togo Marshall, CRI-Enapa, Jasmine 85, CRI-Amankwatia and CRI-Agra rice respectively (Figure 8B). Although, dry biomass increased with increasing nitrogen rates, for Togo Marshall, dry biomass at 0 Kg N/ha (96.7 g) was statistical similar to 60 kg N/ha (105 g) and 90 kg N/ha (92.5 g). The control, recorded statistically similar dry biomass at 30 kg N/ha among varieties such Jasmine 85, CRI-Agra rice, CRI-Amankwatia and CRI-Enapa (Figure 8B).

### 3.8 Association among the growth, yield and yield related parameters, and biomass

Diverse association was observed in the measured parameters as illustrated in Figure 9. Yield exhibited a positive and strong significant association with dry straw weight (p < 0.01, r = 0.85) but had a negative and strong significant association with fresh straw weight (p < 0.05, r = 0.92) (Figure 9). Similarly, growth parameters such as number of tillers (p < 0.01, r = 0.41), number of panicle (p < 0.01, r = 0.52) and plant height (p < 0.01, r = 0.52) had a strong positive and significant association with 1000-grain weight but a weak and significant positive association with culm length (p < 0.05, r = 0.27) (Figure 9). Days to flowering had a strong negative but insignificant association (p>0.05, r = -0.42) with 1000-grain weight. Number of panicle (p < 0.05, r = -0.76), plant height (p < 0.05, r = -0.67) and number of tillers (p < 0.05, r = -0.71) had a strong negative and significant association with days to flowering (Figure 9).

**Figure 9:**
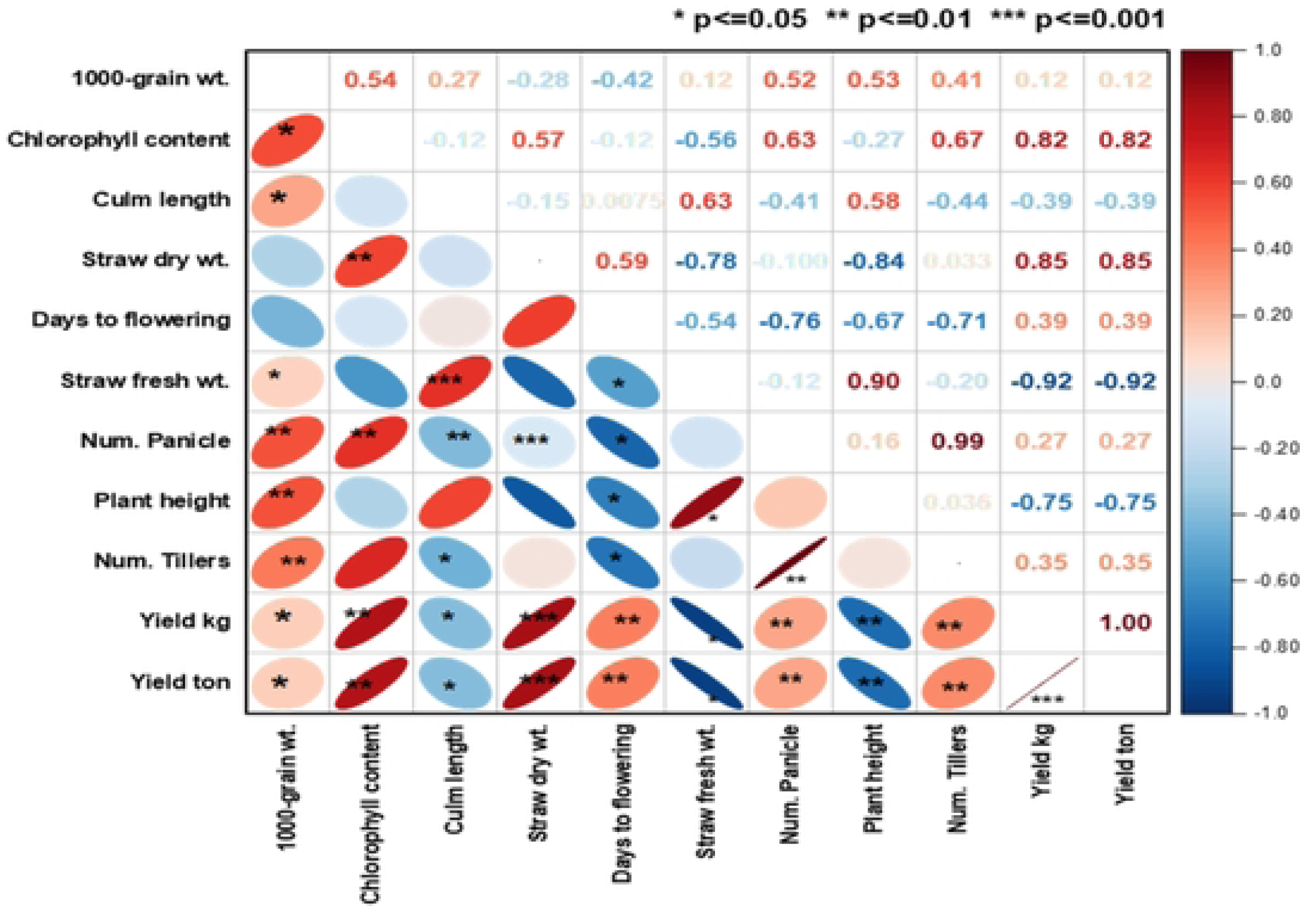
Correlational analysis showing relationship between measured growth and yield traits ofrice cultivated under different nitrogen fertiliser rates. Asterisks indicate significance levels:* p<0.05; P < 0.05P<0.05, ** p<0.01 and*** p<0.001.

## 4.0 Discussion

### 4.1 Variability among the treatment combinations

The study showed that experimental factors including nitrogen application rate, rice variety and season significantly influenced rice growth and yield traits, although the degree of response varied across traits. For plant height and culm length, both variety and nitrogen rate had highly significant effects (p < 0.001), suggesting that genetic makeup and nitrogen availability strongly influence vegetative growth. Singh *et al*. (2017[10]) observed that varietal differences and nitrogen application rates had a highly significant impact on plant height, attributing the variation to genetic potential and nutrient uptake efficiency. Ali *et al*. (2025[11]) reviewed that higher nitrogen rates increased plant height, but the degree of response varied significantly among varieties, reinforcing the role of genetic control over nitrogen responsiveness. Mishra *et al*. (2024[12]) also reported that both genotype and nitrogen application significantly affected culm length, with taller varieties exhibiting greater nitrogen use efficiency under optimal fertilization conditions.

The number of tillers and number of panicles were also significantly influenced by both variety and fertilizer rate (p < 0.001), with a notable interaction between variety and season for tiller number. Similar findings have been reported by Fageria (2007[13]), Reddy *et al.* (2022[14]) and Patel *et al*. (2018[15]) where both variety and fertilizer rate significantly influenced the number of tillers and panicles in rice.

Fresh and dry straw weight were significantly influenced by variety and nitrogen rate (p < 0.001), reflecting enhanced biomass accumulation with higher nitrogen levels. Fageria *et al.* (2003[16]), Singh and Verma (2013[17]) and Ju *et al*. (2021[18]) also reported significant fresh and dry straw weights were in their studies in rice.

For 1000-grain weight, grain yield (kg/ha) and yield (t/ha), nitrogen rate was the major driver (p < 0.001), enhancing yield traits substantially. In line with the current findings, Singh *et al.* (2017[10]) and Mahajan *et al.* (2012[19]) have also reported that nitrogen application rate is a major determinant of 1000-grain weight, grain yield (kg/ha), and overall yield (t/ha) in rice highlighting the critical role of nitrogen in maximizing rice production potential.

### 4.2 Variation among traits in response to nitrogen application

Nitrogen is a vital nutrient for plant growth and development. Nitrogen plays a paramount role in cell growth, elongation, and division of crops (Fathi and Zeidali, 2021[20]). Its deficiency delays phenological development in vegetative and reproductive stages (Fathi and Zeidali, 2021[20]). The results of the present study demonstrated a clear positive correlation between nitrogen application rate and morphological traits of rice. An increase in plant height was observed with increasing N rate particularly at 90 kg N/ha, highlighting the role of nitrogen in cell division, elongation, and overall vegetative growth (Figure 3). Conversely, the substantial reduction in plant height under nitrogen-deficient conditions (0 kg N/ha) further underscores the essential role of nitrogen in plant development (Figure 3). The results corroborate with the reports of Abdou *et al.* (2021[21]) and Mboyerwa *et al*., 2021[22]). Similarly, chlorophyll content of rice increased positively in response to N application with 90 kg N/ha recording the highest chlorophyll content (Figure 3A). Thus, as the nitrogen fertilizer rate increased from 0 kg N/ha to 90 kg N/ha, a consistent increase in chlorophyll content was observed. These findings suggest that increasing nitrogen availability significantly enhances chlorophyll production in the plants. Even so, at lower fertilizer rates, notable improvements in chlorophyll content were observed. The application of 60 kg N/ha resulted in a 25.6% increase in chlorophyll content compared to the control, while 30 kg N/ha led to a 16.5% increase (Figure 3A). This indicates that even moderate levels of nitrogen fertilization can have a meaningful impact on chlorophyll production. The observed relationship between nitrogen fertilization and chlorophyll content is consistent with the fundamental role of nitrogen in chlorophyll synthesis. Nitrogen have been reported as a crucial component of chlorophyll molecules, and its increased availability promotes greater chlorophyll production, which can enhance the plant’s photosynthetic capacity and overall growth potential (Zhou *et al*., 2022[23]). These findings are in line with Iqbal *et al.* (2021[24]) who reported more greenness of leaves at higher N application rate in rice. Swain and Sandip (2010[25]) also reported an increased in SPAD values with an increase in nitrogen levels from 0 to 150 kg N ha^-1^ in rice.

In addition to influencing plant height and chlorophyll content of rice, nitrogen application also significantly affected panicle and tiller production (Figure 3 C-D). The observed increase in the number of panicles and tillers with increasing nitrogen rates indicates nitrogen’s role in promoting reproductive development in rice. The significant reduction in panicle number under 0 N kg/ha, highlights the importance of adequate nitrogen availability for optimal reproductive growth hence highlighting the role of nitrogen in cell elongation and internode expansion. Increased panicle production at higher nitrogen rates is likely due to enhanced tillering and greater nutrient uptake, contributing to improved yield potential. This finding aligns with reports of Zhou *et al.* (2022[23]), Zhou *et al* (2017[26]) and Firouzi (2015[27]) which indicated that, optimized nitrogen fertilizer application (OFA) increases rice yield by improving tiller quality, enhancing panicle development and increasing the number of filled spikelet.

Days to flowering was also significantly affected by nitrogen application (Figure 4.A). Higher nitrogen levels resulted in a delayed flowering period, likely due to the prolonged vegetative growth phase before transitioning to the reproductive stage. Applying the right amount of N fertilizer can significantly increase biomass, and also high biomass is only possible under N fertilization conditions (Fathi *et al*., 2016[28]). The significant positive response of biomass to nitrogen application as demonstrated in this study could be attributed to the role of N to maintain leaf surface survival; as leaf surface durability increases, the duration and rate of leaf photosynthesis also increase, allowing the plant to produce more fresh and dry matter (Fathi, 2022[29]). Nitrogen deficiency stimulates competition for the transfer of this element in the plant, impairs timely and complete formation of reproductive organs by decreasing crop growth rate (CGR), delays plant phenology, lowers harvest index and ultimately reduces grain yield and biomass of plants (Fathi and Zeidali, 2021[20]).

The application of N-fertiliser also showed a direct relationship with yield traits such as grain yield (Kg/ha and t/ha) and 1000-grain weight (Figure 4 A-C). The observed relationship between nitrogen fertilization and yield parameters is consistent with the fundamental role of nitrogen in improving photosynthetic efficiency which in turn enhances overall yield. This trend further suggests that higher nitrogen availability promotes not only vegetative growth but also reproductive development, which is essential for grain yield. Increased 1000-grain weight and grain yield at 90 kg N/ha is likely due to enhanced tillering and panicle formation hence, contributing to improved yield potential. The results align with the work of Shrestha *et al*. (2022[30]), Prasad and Mailapalli (2018[31]) and Jun-li (2014[32]).

### 4.3 Variation among rice varieties in Response to Nitrogen Application

#### 4.3.1 Growth parameters

The photosynthetic apparatus of plants consists mainly of N, a widely used fertilizer in plants (Bassi *et al*., 2018[33]). Hence, N is essential for increasing leaf area, affects plant growth habits and leaf longevity, and ultimately affecting photosynthetic efficiency (Olszewski *et al*., 2014[34]). The revealed significant variation among rice varieties in their response to nitrogen (N) fertilisation (Figure 4 A–E) indicates genotypic differences in nitrogen uptake efficiency, assimilation and utilization (Nguyen *et al.,* 2016[35]). Varieties; Togo Marshall, CRI-Agra rice and Jasmine 85 exhibited over 30% increase in chlorophyll content at 90 kg N/ha indicating the ability of such genotypes to absorb and assimilate nitrogen at highest rate hence leading to enhanced photosynthetic performance (Figure 4.A). Some varieties also require lower fertilisation rate to reach chlorophyll saturation to improve photosynthetic performance. CRI-Enapa had lower chlorophyll content at 90 kg N/ha but produced higher plant height, number of panicles and tiller at 90 kg N/ha compared to other rice varieties in the present study. Responsive rice varieties exhibit greater internode elongation and overall plant height increase due to auxin synthesis under higher fertilisation rates (Shafi *et al*., 2023[36]). Varieties; Togo Marshall and CRI-Agra rice exhibited higher plant height in response to N fertilisation compared to varieties such as CRI-Amankwatia (Figure 4.B). Thus, low-N-responsive varieties may show stunted growth even under higher nitrogen levels, indicating genetic constraints on nitrogen use efficiency (NUE) (Lal *et al*., 2024[37]). Similar varietal differences in nitrogen use efficiency and chlorophyll response have been reported in rice by Peng *et al*. (2021[38]). They reported that, improved varieties recorded higher biomass production with lower chlorophyll content under reduced nitrogen inputs, indicating better nitrogen-use efficiency. Singh and Dwivedi (2016[39]) also observed that rice varieties like ’Swarna’ attained chlorophyll saturation and maximum tillering at moderate nitrogen application.

Increase in number of tillers at highest N fertilisation rate recorded by CRI-Amankwatia, CRI-Enapa and Togo Marshall could have accounted for the increased number of panicles observed among these varieties. Additionally, the positive significant association observed between number of tillers and panicles further justifies the aforementioned observation (Figure 4.C - D). Thus, responsive varieties prioritize converting most tillers into panicle-bearing stems (Mohapatra *et al*., 2025[40]). Varieties with shorter days to flowering have been reported to have relatively smaller leaves and tillers as they transition quickly to the reproductive stage, limiting the time available for tiller formation (Hussien *et al*., 2014[41]; Yan *et al*., 2024[42]). This could have accounted for reduction in panicle production among varieties Jasmine 85 and CR-Agra rice (Figure 4.C).

The observed increase in culm length across all rice varieties in response to nitrogen (N) fertilization suggests that nitrogen plays a crucial role in promoting stem elongation and overall plant height (Figure 4.E). The positive correlation between nitrogen application and culm length can be attributed to several physiological and biochemical mechanisms. Nitrogen is a fundamental component of amino acids, proteins and enzymes involved in cell division and elongation (Luo *et al*., 2020[43]). Higher nitrogen availability enhances the synthesis of structural proteins and growth regulators such as gibberellins, which promote internode elongation and contribute to increased culm length (Zimmermann *et al*., 2021[44]). This explains why culm length increased progressively with nitrogen application across all varieties (Figure 4.E). The differential responses among varieties highlight genetic variability in nitrogen use efficiency (NUE) and sensitivity to nitrogen-driven growth promotion. CRI-Agra rice, which exhibited the longest culm length at 90 kg/ha, may have a higher capacity to assimilate and utilize nitrogen for stem elongation. Conversely, Jasmine 85, which recorded the least increase in culm length, may have either lower NUE or genetic constraints in internode elongation, limiting its responsiveness to nitrogen application. Sun *et al*. (2020[45]) and Pan *et al.* (2019[46]) reported similar results in rice. Furthermore, the significant increase in culm length in Togo Marshall, despite having the shortest initial culm length at 0 kg/ha, suggests that this variety exhibits a high nitrogen responsiveness under fertilized conditions (Figure 4.E). This could be attributed to a greater plasticity in internode elongation when nitrogen availability is improved. The relatively lower response in Jasmine 85 could be due to a genetically determined shorter culm, where the variety prioritizes resource allocation towards reproductive rather than vegetative growth.

#### 4.3.2 Biomass traits

The significant variation in straw fresh weight and dry straw weight among rice varieties in response to nitrogen (N) fertilization highlights the differential nitrogen use efficiency (NUE) and biomass accumulation potential of each variety (Figure 5 A - B). The observed trends suggest that genetic factors, physiological traits and nitrogen uptake efficiency play critical roles in determining biomass responses to nitrogen application. Rice varieties differ in their ability to uptake, assimilate and utilize nitrogen, which influences their biomass production. The higher fresh weight response in CRI-Enapa and Togo Marshall at 90 kg N/ha suggests that these varieties have a greater nitrogen uptake efficiency (NUpE) and higher nitrogen utilization efficiency (NUtE) under increased nitrogen availability. This aligns with studies indicating that high-N-responsive rice genotypes often exhibit superior root system architecture, enhanced nitrate reductase activity and efficient nitrogen remobilization to support biomass accumulation (Hoyt, 2022[47]; Bharati *et al*., 2020[48]; Bharati *et al*., 2019[49]). Conversely, CRI-Amankwatia, which exhibited the lowest increase in fresh straw weight, may have a lower capacity for nitrogen uptake or a limited ability to convert absorbed nitrogen into biomass (Figure 5A). This could be due to differences in root morphology, nitrogen transporters or internal nitrogen allocation patterns. The increase in biomass with nitrogen application can also be linked to enhanced photosynthesis due to enhanced chlorophyll content. Nitrogen is a key component of chlorophyll and rubisco, the enzyme responsible for carbon fixation. CRI-Enapa and Togo Marshall, which recorded higher biomass accumulation at 90 kg N/ha had higher leaf chlorophyll content, increased light interception and greater photosynthetic efficiency, leading to improved carbon assimilation and dry matter production (Figure 4A). The differences in nitrogen response among varieties may also be influenced by how nitrogen is partitioned between vegetative and reproductive organs. Varieties, CRI-Amankwatia and CRI-Agra rice, which showed similar fresh straw weight at 0 kg N/ha, 30 kg N/ha, and 60 kg N/ha, might allocate nitrogen preferentially to reproductive structures rather than vegetative biomass. This trade-off between biomass and reproductive growth has been reported in nitrogen-efficient rice varieties by Srikanth *et al*. (2023[3]); Wang *et al*. (2022[4]) and Liu *et al.* (2015[49])

#### 4.3.3 Yield traits

The observed delay in flowering across all rice varieties with increasing nitrogen application suggests that nitrogen availability plays a crucial role in regulating the transition from the vegetative to the reproductive phase (Luo *et al*., 2020[43]). Nitrogen is known to promote vegetative growth by enhancing cell division, chlorophyll content, and photosynthetic efficiency (Pérez-Álvarez *et al*., 2024[50]). Consequently, excess nitrogen may prolong the vegetative phase, delaying the expression of floral genes such as heading date 3a (Hd3a) and rice flowering locus T1 (RFT1), which are critical for floral induction (Tamaki *et al*., 2007[51]). The extent of this delay varied among varieties, likely due to genetic differences in nitrogen use efficiency (NUE) and sensitivity to nitrogen-induced hormonal regulation. Togo Marshall and Jasmine85 exhibited the most pronounced delay, possibly due to their higher responsiveness to nitrogen in terms of biomass accumulation before flowering (Figure 5A).

The improvement in 1000-grain weight with increasing nitrogen levels indicates the role of nitrogen in grain filling and assimilate translocation. Nitrogen enhances the synthesis of proteins and starch, which are essential for grain development (Peng *et al*., 2014[52]). The variation in grain weight response across varieties could be attributed to differences in sink strength, where some varieties (CRI-Enapa) may have a greater ability to accumulate and translocate photosynthates into grains compared to others. The significantly higher grain weight in CRI-Enapa suggests superior nitrogen partitioning, which enhances grain filling duration and grain size (Figure 5C).

Rice yield improvement with nitrogen application can be explained by increased tillering, enhanced leaf area index and prolonged photosynthetic activity (Fageria and Baligar, 2005[53]). However, the magnitude of yield increase varied among varieties, likely due to differences in nitrogen uptake efficiency and source-sink dynamics. The highest yield recorded in Togo Marshall at 90 kg/ha suggests a strong balance between vegetative growth and reproductive output. In contrast, CRI-Enapa exhibited the highest percentage increase in yield (61.63%), indicating a higher responsiveness to nitrogen fertilization (Figure 5C). This could be linked to enhanced nitrogen uptake efficiency, improved biomass accumulation and efficient nitrogen remobilization to reproductive organs. Mingotte *et al*. (2013[54]) and Hawkesford (2017[55]) reported similar results in their studies. Genetic variability plays a crucial role in N response, with certain cultivars consistently outperforming others across N rates. These findings emphasize the importance of variety-specific N management strategies to optimize yield while avoiding excessive fertilizer use (Jahan *et al*., 2022[56]).

The yield observed in CRI-Agra Rice, CRI-Amankwatia, and Togo Marshall under different nitrogen levels suggests that these varieties maintain a relatively efficient nitrogen use strategy. The moderate increase in Jasmine85, despite its lower grain weight, implies that it may have limitations in nitrogen assimilation and grain filling efficiency. Excessive nitrogen application can sometimes lead to luxury consumption, where the plant takes up more nitrogen than needed, potentially leading to lodging, excessive vegetative growth, and reduced harvest index (Wei *et al*., 2023[57]).

Overall, the interaction between nitrogen application and rice variety influences key agronomic traits, primarily through nitrogen’s role in modulating vegetative and reproductive growth. Understanding these mechanisms will be crucial for optimizing nitrogen management strategies tailored to specific rice varieties, ensuring both productivity gains and environmental sustainability

### 4.4 Relationship between growth, biomass and yield and yield related parameters

The observed associations among yield, biomass and morphophysiological traits suggest that key agronomic parameters play a crucial role in determining rice productivity (Figure 9). The strong positive correlation between yield and dry straw weight implies that higher biomass accumulation contributes positively to grain production. This relationship may be attributed to enhanced photosynthetic efficiency and resource allocation, where dry matter is effectively translocated from vegetative structures to reproductive organs. Conversely, the negative and strong correlation between yield and fresh straw weight suggests that excessive vegetative growth in terms of fresh biomass may not necessarily translate to higher grain yield. This could be due to excessive water retention in fresh biomass, which may compete with grain filling for available assimilates.

The significant positive correlations of number of tillers, number of panicles and plant height with 1000-grain weight suggest that these traits contribute to overall grain development and yield stability (Figure 9). Higher tillering capacity and panicle production likely enhance the grain sink potential, leading to increased grain weight (Parida *et al*., 2022[58]). Additionally, taller plants might have better light interception and photosynthetic activity, which could further promote grain filling.

The negative but insignificant correlation between days to flowering and 1000-grain weight (Figure 9) suggests that late-flowering varieties might not necessarily produce heavier grains. The weak association might indicate that while early-flowering plants allocate more resources toward grain filling, late-flowering varieties might experience environmental stressors (e.g., drought or high temperatures) that limit their grain-filling capacity (Wingler and Soualiou, 2025[59]; Chen *et al*., 2023[60]). Moreover, the strong negative and significant associations between days to flowering and number of panicles, plant height and number of tillers further emphasize the trade-offs between vegetative growth duration and reproductive output. Early-flowering plants may prioritize reproductive development over excessive vegetative growth which may explain why they produce fewer tillers and panicles but potentially higher grain yield efficiency (Olliff-Yang *et al*., 2021[61]; Hossain *et al*., 2024[62]). The trade-off between early flowering and vegetative biomass has been reported as a drought escape mechanism in cereals such as rice maize, wheat and barley (Jagadish *et al*., 2012[63]; Xi et al., 2023[64]).

## 5.0 Conclusion

The findings of the study underscore the significant influence of nitrogen application on the morpho-agronomic, biomass and yield parameters of rice genotypes. Nitrogen, being a critical nutrient for plant growth and development, positively correlated with key growth traits such as plant height, chlorophyll content, tiller production, panicle number, and overall grain yield. The observed variations among rice genotypes in their response to nitrogen fertilization highlight the importance of genotype-specific nitrogen management strategies to optimize productivity and resource use efficiency. Varieties; CRI-Enapa and Togo Marshall exhibited superior nitrogen uptake and utilization efficiency compared CRI-Amankwatia and Jasmine 85. The observed trends indicate that while higher nitrogen levels generally enhance biomass accumulation and yield, the efficiency of nitrogen use varies across genotypes. Also, the significant relationship between nitrogen application and yield, biomass and growth parameters reinforce that optimized nitrogen fertilization not only improves yield but also influences the physiological and biochemical processes essential for rice growth.

Genotypes like Togo Marshall and CRI-Agra rice which demonstrated a strong response to nitrogen fertilization, are suitable candidates for high-input farming systems. Optimizing nitrogen use will be key to increasing rice yields while minimizing environmental impacts such as nitrogen leaching and greenhouse gas emissions.

## Acknowledgements

We thank CSIR-Crops Research Institute for their assistance and Senior members and field staff of CRI rice Improvement program.

## Author contributions

**Boadu Sober Ernest** was involved in the conceptualization, data curation, methodology, investigation and writing of the original draft of the manuscript.

**Esther Fobi Donkor, Charles Afriyie-Debrah and Maxwell Darko Asante** were involved in the conceptualization, supervision, formal analysis and reviewing and editing of the manuscript.

Ralph K. Bam, Priscilla Francisco Ribeiro, Kirpal Agyemang Ofosu, Daniel Dzorkpe Gamenyah and Vincent Opoku Agyemang were involved in the data curation, investigation and methodology of the research.

**Samuel Novo** was involved in the reviewing and editing of the manuscript and formal analysis

